# Prediction of occult tumor progression via platelet RNAs in a mouse melanoma model: a potential new platform of cancer screening for early detection of cancer

**DOI:** 10.1101/2021.02.09.430530

**Authors:** Yue Yin, Ruilan Jiang, Mingwang Shen, Zhaofang Li, Ni Yan, Junqiao Feng, Hong Jiang, Jiaxin Lv, Lijuan Shi, Lina Wang, Xi Liu, Kaiyun Zhang, Di Chen

**Author notes:** Materials & Correspondence Correspondence and material requests should be addressed to Dr. Yue Yin.

## Abstract

Cancer screening provides the opportunity to detect cancer early, ideally before symptom onset and metastasis, and offers an increased opportunity for a better prognosis. The ideal biomarkers for cancer screening should discriminate individuals who have not developed invasive cancer yet but are destined to do so from healthy subjects^1,2^. However, most cancers lack effective screening recommendations. Therefore, further studies on novel screening strategies are urgently required. Here, our proof-of-concept study shows blood platelets could be a platform for liquid biopsy-based early cancer detection. By using a simple suboptimal inoculation melanoma mouse model, we identified differentially expressed RNAs in platelet signatures of mice injected with a suboptimal number of cancer cells (eDEGs) compared with mice with macroscopic melanomas and negative controls. These RNAs were strongly enriched in pathways related to immune response and regulation. Moreover, 36 genes selected from the eDEGs via bioinformatics analyses were verified in a mouse validation cohort via quantitative real-time PCR. LASSO regression was employed to generate the prediction models with gene expression signatures as the best predictors for occult tumor progression in mice. The prediction models showed great diagnostic efficacy and predictive value in our murine validation cohort, and could discriminate mice with occult tumors from control group (area under curve (AUC) of 0.935 (training data) and 0.912 (testing data)) (gene signature including *Cd19, Cdkn1a, S100a9, Tap1*, and *Tnfrsf1b*) and also from macroscopic tumor group (AUC of 0.920 (training data) and 0.936 (testing data)) (gene signature including *Ccr7, Cd4, Kmt2d*, and *Ly6e*). Our study provides evidence for potential clinical relevance of blood platelets as a platform for liquid biopsy-based early detection of cancer. Furthermore, the eDEGs are mostly immune-related, not tumor-specific. Hence it is possible platelets-based liquid biopsy could enable simultaneous early detection of cancers from multiple organs of origin^3^. It is also feasible to determine the origin of cancer since platelet profiles are influenced by tumor type^3^.

## Main text

Traditional cancer screening methods have demonstrated low accuracy and efficacy, while novel cancer markers such as circulating tumor cells (CTCs) and circulating tumor DNA (ctDNA), which offer new genomic approaches through liquid biopsies, still have limited efficiency^1,4–6^. The ideal biomarkers for the early detection of cancer should discriminate individuals with occult cancer that is destined to progress from healthy subjects. Therefore, further studies searching for new blood-based biomarkers for early cancer detection are now urgently required.

Blood platelets, which are traditionally known for their function in hemostasis and thrombosis, have emerged as important participants in tumor pathogenesis^7,8^. Recent studies have indicated significant platelet involvement in cancer growth and metastasis^9,10^. It has been reported that tumor-educated platelets (TEPs) may have potential for cancer companion diagnostics^3,11–13^. However, whether platelets could serve as a platform for cancer risk assessment or early disease diagnostics still merits further investigation.

### Inoculation of suboptimal numbers of tumor cells can induce delayed melanoma formation in mice

To investigate these issues, we established a melanoma mouse model by inoculating C57BL/6 mice with a suboptimal number of tumor cells, which were named an “early-early” mouse model since the tumors in our model were occult or microscopic before they rapidly grew and became macroscopic. C57BL/6 mice were subcutaneously injected with several concentrations of B16F10 cells. In contrast with the rapid tumor development in mice from the group injected with a optimal number of 1 × 10^5^ cells per mouse established by previous studies^14,15^, inoculation of mice with lower numbers of cells postponed the onset of tumor formation, with variable growth kinetics, as evidenced by the dispersion of growth curves (Fig. 1a). In groups injected with 1 × 10^5^ cells and 1 × 10^4^ cells per mouse, all mice developed tumors which became visible in 2 weeks and 3 weeks post-inoculation respectively (Fig. 1a, b). However, injection with suboptimal numbers of cells (5 × 10^3^ cells or 2 × 10^3^ cells per mouse) induced tumors in some subjects that remained small and did not progress for as long as 6 weeks after inoculation (Fig. 1a, b). Around 76% of mice (100 out of 131) injected with 5 × 10^3^ cells per mouse developed tumors that became visible at 2-6 weeks after inoculation, while only 13% of mice (8 out of 60) injected with 2 × 10^3^ cells per mouse formed visible tumors within 6 weeks post-inoculation (Fig. 1b, c). Moreover, around 24% of mice from the group injected with 5 × 10^3^ cells per mouse and 87% of mice injected with 2 × 10^3^ cells did not develop melanomas within 6 weeks after inoculation and remained tumor-free for a prolonged period of 15 weeks post-inoculation (Fig. 1b, c).

**Fig. 1.**
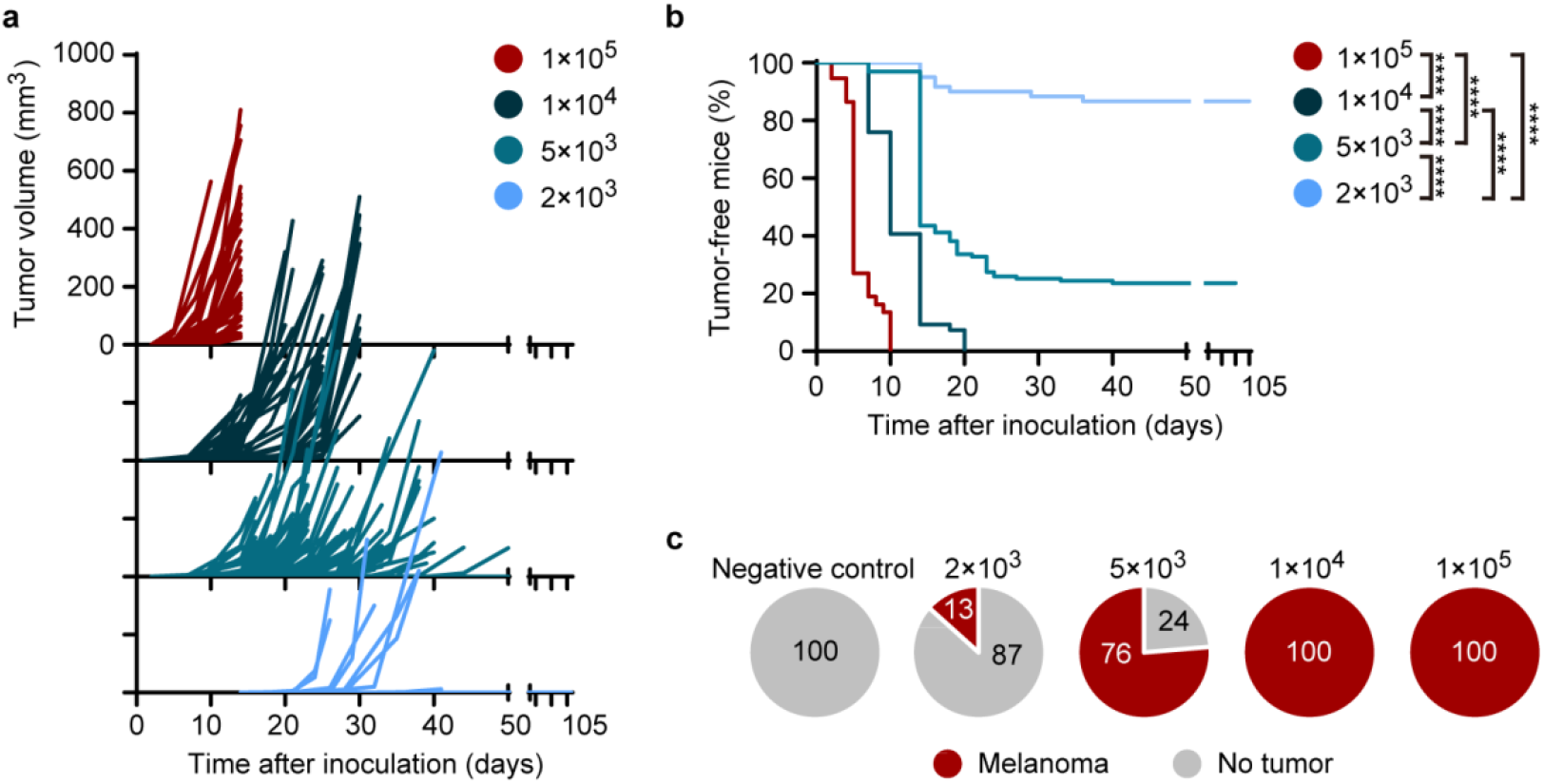
Tumor growth kinetics in C57BL/6 mice inoculated with different numbers of B16F10 cells. **a**, Tumor growth curves after inoculation with optimal and suboptimal numbers of B16F10 cells. **b, c**, Proportion of tumor-free and tumor-bearing mice following inoculation with B16F10 cells. Data pooled from *n* = 8 (1 × 10^5^ cells per mouse, red), *n* = 12 (1 × 10^4^ cells per mouse, dark blue), *n* = 29 (5 × 10^3^ cells per mouse, blue) and *n* = 14 (2 × 10^3^ cells per mouse, light blue) biologically independent experiments with *n* = 37 mice (1 × 10^5^ cells per mouse), *n* = 54 mice (1 × 10^4^ cells per mouse), *n* = 131 mice (5 × 10^3^ cells per mouse) and *n* = 60 mice (2 × 10^3^ cells per mouse). \*\*\*\**P* < 10^−10^, log-rank Mantel-Cox test (**b**).

Since some tumors developed late (Fig. 1a), these “late-developer” mice harbored occult or microscopic melanomas after inoculation for weeks before they progressed into macroscopic tumors afterwards. We proposed that the “pre-diagnostic” blood samples from mice inoculated with a suboptimal number of tumor cells could be used for screening novel “early-early” cancer biomarkers.

### Platelet mRNA profiles of mice inoculated with a suboptimal number of tumor cells are distinct from those of both healthy and tumor-bearing mice

To screen for novel “early-early” cancer biomarkers in our melanoma model, we collected blood samples from optimal inoculation group (1 × 10^5^ cells per mouse, O group), suboptimal inoculation group (2 × 10^3^ cells per mouse, S group), and negative control group (C group) on day 14 post-injection, when optimal inoculation group all developed palpable tumors and suboptimal inoculation group did not form visible tumors yet, indicating the tumors might still be occult (Fig. 2a). Mice were autopsied after blood collection and examined under a magnifying scope to confirm there were no visible tumors in S group (Extended Data Fig. 1a).

**Fig. 2.**
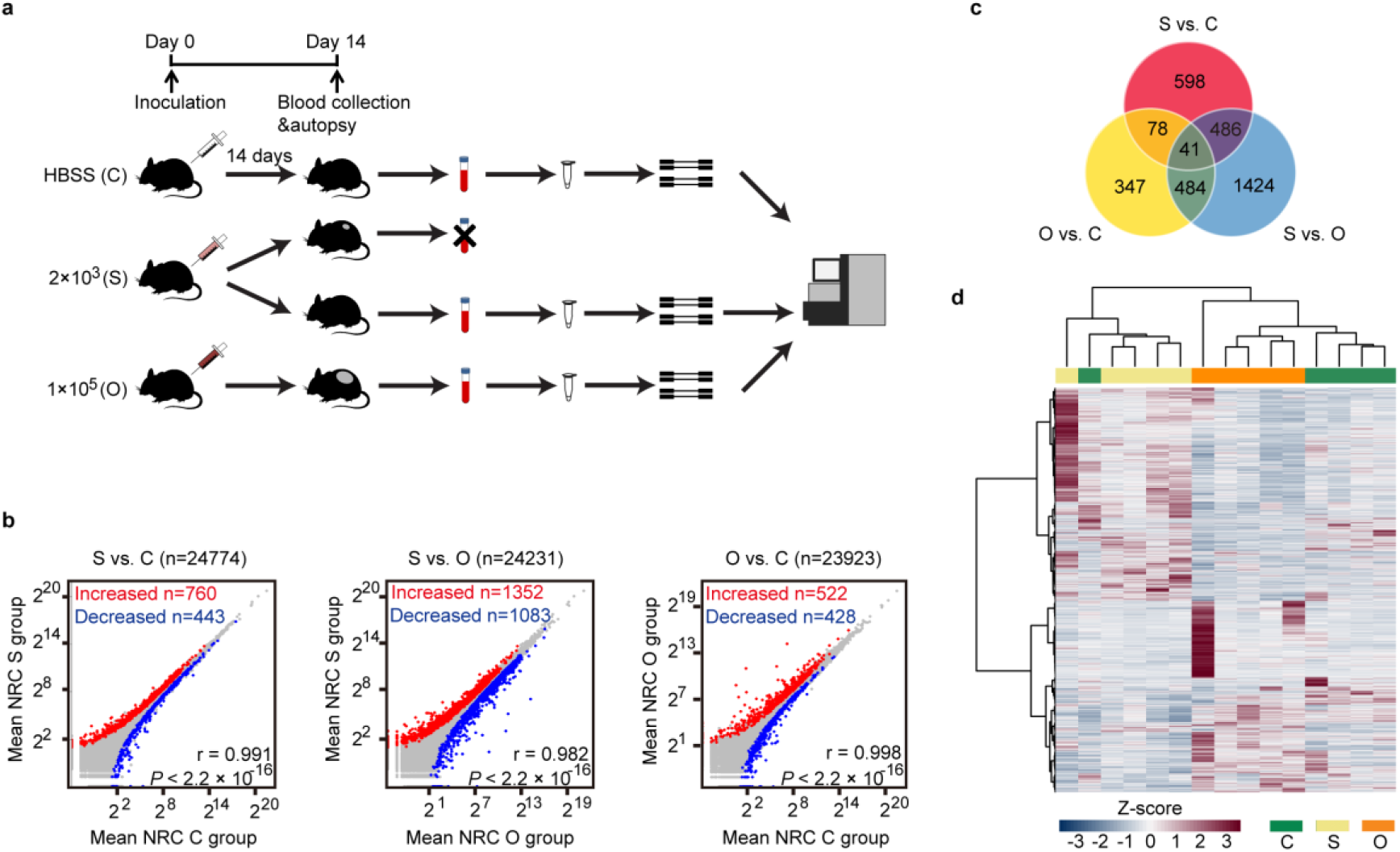
Platelet RNA profiles of mice inoculated with a suboptimal number of B16F10 cells are distinct from those of control mice and mice with macroscopic tumors. **a**, Animal model and platelet mRNA sequencing workflow, as starting from B16F10 cell injection, terminal blood collection, to platelet isolation, and mRNA sequencing. C group: negative control group, mice injected with HBSS (Hank’s Balanced Salt Solution); S group: suboptimal inoculation group, mice inoculated with 2 × 10^3^ B16F10 cells; O group, optimal inoculation group, mice inoculated with 1 × 10^5^ B16F10 cells (**a**-**d**). **b**, Correlation plots of mRNAs detected in platelets of S group, C group and O group mice, including highlighted increased (red) and decreased (blue) platelet mRNAs. NRC, normalized read counts (mean of group). r value calculated from Pearson’s correlation test. **c**, Venn diagram of differentially expressed genes from pairwise comparisons. **d**, Heatmap of hierarchical clustering of platelet mRNA profiles of S group (beige), C group (green) and O group (orange). Data pooled (**b**-**c**) from *n* = 5 biologically independent experiments with *n* = 50 (S group) or *n* = 40 (C group) and *n* = 20 (O group) mice, or data representing all 5 independent experiments (**d**).

Recently, it has been reported that tumor-educated platelets (TEPs) may have potential for cancer companion diagnostics^3,11–13^. Therefore, we isolated blood platelets for further study as well as peripheral blood mononuclear cells (PBMCs) for comparison. Both platelet RNA and PBMC RNA were isolated and evaluated for quantity and quality. Total platelet RNA samples of 20 mice in O group, 50 mice in S group and 40 mice in C group were pooled into 5 samples per group in order to meet the quantity criteria of RNA sequencing. The disparity in platelet RNA quantity of mice from different groups was probably due to higher platelet production via thrombocytosis in tumor-bearing mice^16^. Total PBMC RNA samples of 20 mice in O group, 24 in S group and 25 in C group were also pooled into 5 samples per group to guarantee sufficient RNA quantity. Pooled platelet and PBMC RNA samples were then processed for RNA sequencing. Platelet RNA sequencing yielded a mean read count of around 53 million clean reads per sample, while PBMC RNA sequencing yielded about 44 million clean reads per sample. After genome mapping of RNA reads, we identified among the platelet RNAs known platelet-abundant genes, such as *B2m* (beta-2 microglobulin), *Fth1* (ferritin heavy polypeptide 1), *Pf4* (platelet factor 4), *Ppbp* (pro-platelet basic protein) and *Tmsb4x* (thymosin, beta 4, X chromosome) (Extended Data Fig. 1b), which yielded much higher read counts than average level. The obtained platelet RNA profiles correlated with PBMC RNA profiles, but the correlation between platelet and PBMC RNA profiles was much less prominent than that between samples within the platelet or PBMC group (Extended Data Fig. 1c). Moreover, mRNAs such as *B2m, Tmsb4x, Ppbp, Rgs18* (regulator of G-protein signaling 18), *Ctsb* (cathepsin B), *Calr* (calreticulin), *Eef2* (eukaryotic translation elongation factor 2) and *Igfbp4* (insulin-like growth factor binding protein 4), which were previously reported differentially expressed genes between platelet and PBMC profiles^17^, were also differentially expressed in our sequencing data (Extended Data Fig. 1d).

A total of 760 out of 24,774 mRNAs were increased and 443 out of 24,774 mRNAs were decreased in platelet samples of S group as compared to samples of C group, while presenting a strong correlation between these platelet profiles (r = 0.991, Pearson’s correlation) (Fig. 2b left). A total of 1,352 out of 24,231 mRNAs were increased and 1,083 mRNAs were decreased in S group compared with O group (r = 0.982, Pearson’s correlation) (Fig. 2b middle). Out of 23,923 mRNAs, 522 were increased and 428 were decreased in O group compared to C group (r = 0.998, Pearson’s correlation) (Fig. 2b right). For PBMC samples, a total of 239 out of 31,625 mRNAs were increased and 251 out of 31,625 mRNAs were decreased in PBMC samples of S group as compared to samples of C group, also presenting a strong correlation between these PBMC mRNA profiles (r = 0.996, Pearson’s correlation) (Extended Data Fig. 2a left). A total of 732 out of 31,263 mRNAs were increased and 1,477 mRNAs were decreased in S group compared with O group (r = 0.833, Pearson’s correlation) (Extended Data Fig. 2a middle). Out of 26,630 mRNAs, 1,676 were increased and 868 were decreased in O group compared to C group (r = 0.803, Pearson’s correlation) (Extended Data Fig. 2a right). We detected in platelets 3,458 and in PBMCs 3,694 differentially expressed protein coding and non-coding RNAs by multiple pairwise comparisons, which were used for subsequent investigations (Fig. 2c, Extended Data Fig. 2b). Hierarchical clustering based on differentially detected platelet mRNAs distinguished 3 sample groups with minor overlap, while clustering based on PBMC mRNAs could not quite discriminate S group from C group (Fig. 2d, Extended Data Fig. 2c).

### Blood platelets provide novel biomarkers to predict occult tumor progression in mice

To screen for distinct markers of occult melanoma, we searched for genes differentially expressed in S group compared with both C group and O group (eDEGs, “e” as in “early-early mouse model”) (Fig. 3a, details in “Methods”). Compared with 524 eDEGs (436 were protein-coding genes) from platelet data, there were only 149 genes (only 63 were protein-coding genes) in PBMC data that fulfilled our criteria (Supplementary Table 1, 2). KEGG pathway analysis revealed that these differentially expressed mRNAs from platelets of suboptimal inoculation group (eDEGs) were strongly enriched for biological processes related with immune response or regulation, such as “cytokine receptor interaction” and “cell adhesion molecules” (Fig. 3b, Extended Data Table 1), whereas the differentially expressed genes in PBMC mRNAs were only enriched for two biological processes with low gene counts in each pathway (Extended Data Fig. 2d, Extended Data Table 2). Therefore, platelet RNA profiles might provide a platform for screening novel biomarkers of occult tumor. Furthermore, platelets were the optimum biosource for screening new biomarkers for early cancer detection since PBMC RNA profiles failed to yield enough eDEGs or significantly enriched pathways. To better understand the interplay among the identified eDEGs in platelets, we obtained the protein-protein interaction (PPI) network using the online STRING tool^18^. The complicated network was made up of 25 modules, including 351 nodes and 1,148 edges and the top three significant modules were selected for further analysis (Fig. 3c-e). The first module contained 26 nodes and 270 edges, including *Tlr9* (toll-like receptor 9), *Icam1* (intercellular adhesion molecule 1), *Ccr7* (chemokine (C-C motif) receptor 7), *Ifng* (interferon gamma), etc (Fig. 3c). In the second and third module, there were only 10 nodes found and the degree values of genes (edges) were much lower than those in the first module (Fig. 3d, e). Combining literature search with our bioinformatics analyses, we finally selected 36 genes from our platelet data for subsequent experimental validation (Fig. 3f).

**Fig. 3.**
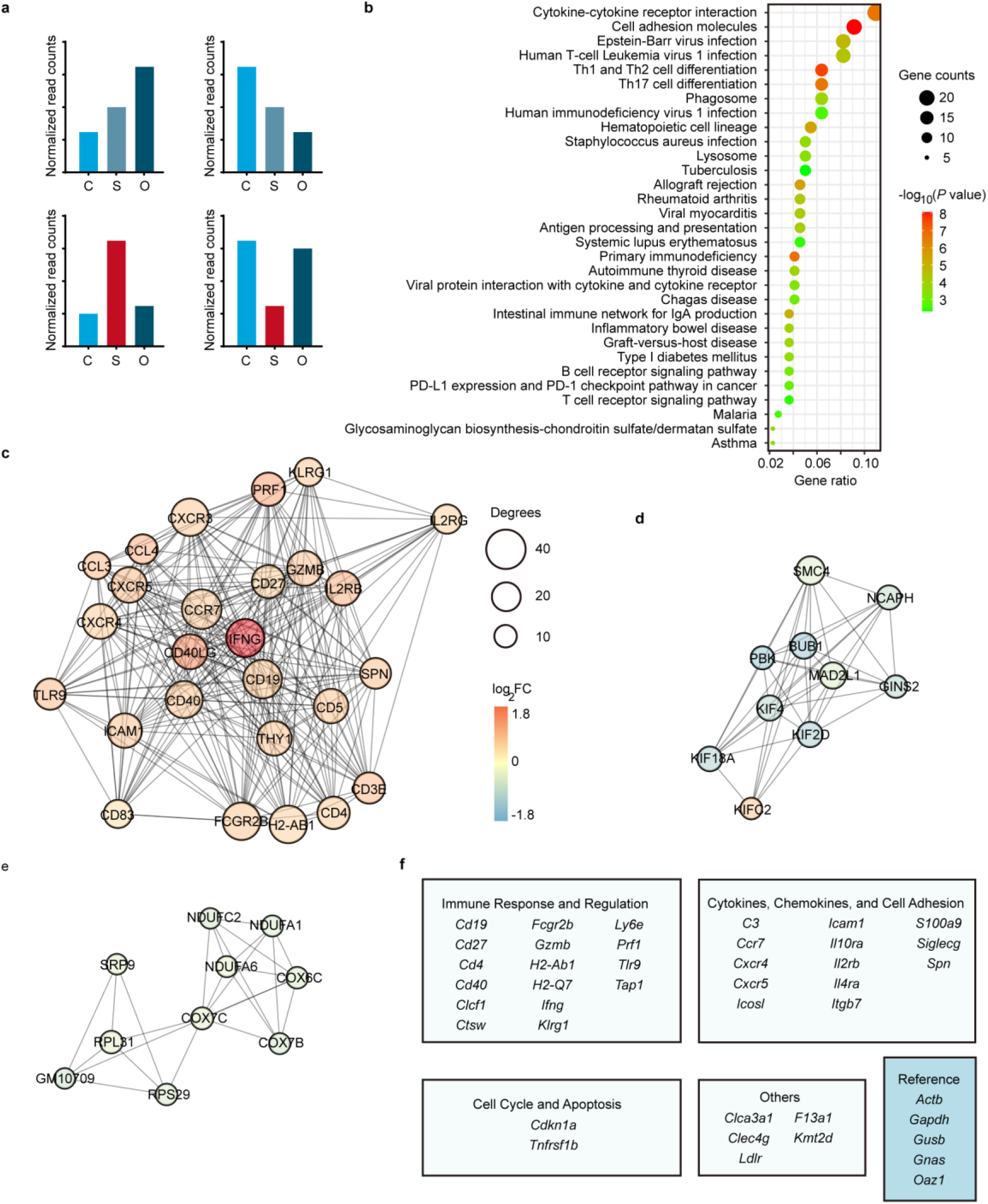
Bioinformatics analyses of differentially expressed genes from suboptimal inoculation group. **a**, Schematics of screening strategy for differentially expressed genes for screening early cancer biomarkers (eDEGs). Eligible genes differentially expressed between S (suboptimal inoculation group, mice inoculated with 2 × 10^3^ cells) and C group (negative control group, mice injected with HBSS), and also between S and O group (optimal inoculation group, mice inoculated with 1 × 10^5^ cells), shown in red columns (eligible eDEGs criteria see details in “Methods”). **b**, Top GO terms of pathway enrichment analysis of eligible eDEGs with reference from KEGG pathways. Adjusted *P* value < 0.05, Benjamini and Hochberg method. **c**-**e**, Top 3 PPI networks modules of eligible eDEGs. Color of a node in the PPI network: log_2_ (Fold change, FC) value of normalized read counts of genes from S group compared with C group; Size of a node: number of interacting proteins with the designated protein (**c**-**e**). **f**, Panel of 36 genes screened from mRNA sequencing data besides 5 reference genes.

We established a validation cohort using our “early-early” melanoma model to test the diagnostic efficacy and predictive value of the aforementioned 36 biomarkers. Mice inoculated in our previous experiments were divided into 3 groups according to their tumor development status on the day of blood collection (day 14 post-inoculation) (Fig. 4a). Mice with occult tumors on day 14 post-inoculation, which were validated by subsequent tumor progression at least 5 days after blood collection, were categorized as early-early tumor group (E group) (Extended Data Fig. 3, Extended Data Table 3, details in “Methods”). Mice with macroscopic tumors were classified as melanoma group (M group) and mice injected with HBSS were in negative control group (C group) (Fig. 4a). Tumor volumes measured on blood collection day showed that E group mice had no visible tumor or only small tumors (volume < 1 mm^3^) that did not progress for more than 2 weeks post-inoculation, verified by multi-phase regression analysis via Joinpoint program (Fig. 4b, Extended Data Fig. 3, Extended Data Table 3)^19^. Tumor growth kinetics demonstrated that E group mice eventually developed macroscopic melanomas afterwards, much later than M group (Fig. 4c, d).

**Fig. 4.**
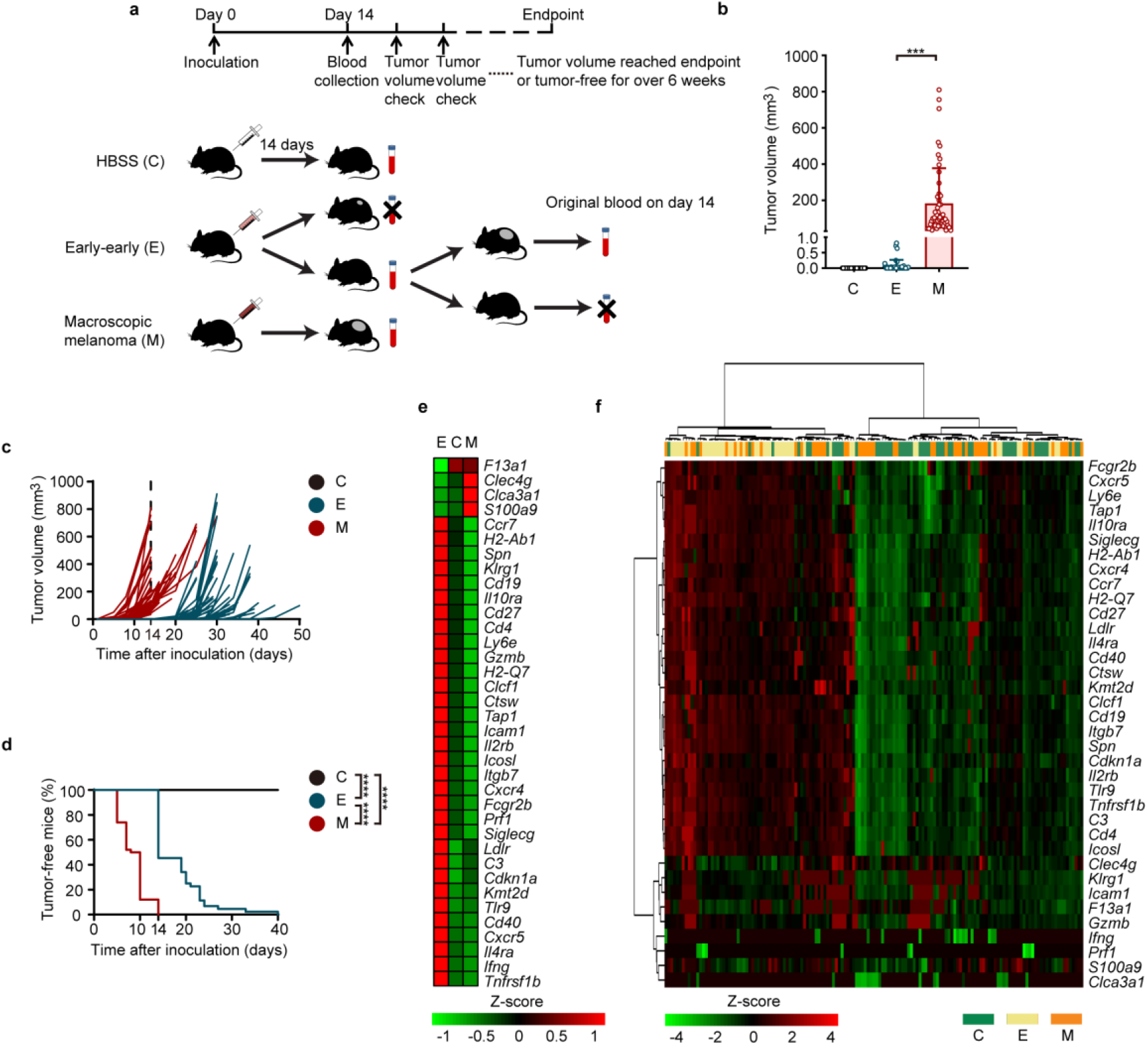
Validation of the expression levels of selected 36 genes via quantitative real-time PCR in a mouse cohort. **a**, The construction workflow of a murine validation cohort, as starting from B16F10 cell inoculation, nonterminal blood collection, to platelet isolation, and observation of tumor developments. C group: negative control group, mice injected with HBSS (Hank’s Balanced Salt Solution); E group: early-early group, mice with occult tumors 14 days post-injection (details in “Methods”); M group, mice with macroscopic tumors 14 days post-injection (**a**-**f**). **b**, Tumor volumes of three groups of mice 14 days after inoculation. \*\*\**P* < 0.001, Kruskal-Wallis non-parametric test, Error bars representing SD values. **c**, Tumor growth kinetics of three groups from the mouse cohort. **d**, Proportion of tumor-free and tumor-bearing mice in three groups of the mouse cohort. \*\*\*\**P* < 10^−10^, log-rank Mantel-Cox test. **e, f**, Heatmap of hierarchical clustering of expression levels of 36 selected genes from platelet mRNA sequencing data (normalized read counts) (**e**) or from quantitative real-time PCR results (Ct_Ref_ - Ct_Gene_) (**f**) in C group (green), E group (beige) and M group (orange). QPCR Data pooled from *n* = 11 biologically independent experiments with *n* = 51 (C group, black), *n* = 44 (E group, blue), and *n* = 50 (M group, red) mice (**b**-**d, f**).

Quantitative real-time PCR (qPCR) experiments were performed to validate the selected 36 genes from eDEGs in our mouse validation cohort (Extended Data Table 4). The normalized expression levels of the 36 genes were mostly in accordance with our previous sequencing data, except *Clca3a1* (chloride channel accessory 3A1), *F13a1* (coagulation factor XIII, A1 subunit), *Ifng, Prf1* (perforin 1 (pore forming protein)) and *S100a9* (S100 calcium binding protein A9), which yielded non-significant results (Extended Data Fig. 4, Extended Data Table 5). Hierarchical clustering of ΔCt values could discriminate E group from C group and M group, which was consistent with our sequencing data (Fig. 4e, f). Although most selected genes could be validated in qPCR experiments, the quantities of several platelet RNA samples were too low for 36-gene-panel qPCR experiments. Therefore, 10 samples were excluded for subsequent analysis. Moreover, some markers such as *Clca3a1, Ifng* or *Prf1*, yielded invalid Ct value in over 20% of samples from each group, probably due to low abundance in platelets (Extended Data Table 5). Therefore, 7 genes including *Clca3a1, F13a1, Gzmb* (granzyme B), *Icam1, Ifng, Klrg1* (killer cell lectin-like receptor subfamily G, member 1), and *Prf1*, were not used for subsequent regression analysis of E and C group. However, *Gzmb* was included in regression analysis of E and M group since it yielded valid Ct results in more than 90% of samples in each group (Extended Data Table 5). Ultimately there were 29 genes and 30 genes included as independent variables in subsequent regression analyses of E vs. C group and E vs. M group.

LASSO binomial logistic regression was applied to generate the prediction model with a multi-gene expression signature as the best predictor for occult tumor progression in mice. Cross-validation was carried out in 10 folds to prevent overfitting (internal training sets and internal validation sets constructed randomly) (Extended Data Fig. 5a, b). Finally the optimal gene signature consisting of *Cd19* (CD19 antigen), *Cdkn1a* (cyclin-dependent kinase inhibitor 1A (P21)), *S100a9, Tap1* (transporter 1, ATP-binding cassette, sub-family B (MDR/TAP)), *Tnfrsf1b* (tumor necrosis factor receptor superfamily, member 1b) for E vs. C group, and *Ccr7, Cd4* (CD4 antigen), *Kmt2d* (lysine (K)-specific methyltransferase 2D), *Ly6e* (lymphocyte antigen 6 complex, locus E) for E vs. M group, as well as the corresponding coefficients were identified by the regularization process of LASSO regression (Extended Data Fig. 5c). Predictive scores for tumor progression were calculated from qPCR data using the training set of 63 mice for E vs. C group and 60 mice for E vs. M group. The scores were then tested in the validation set of 27 and 25 mice for E vs. C and for E vs. M group respectively. The biomarker score formula for E vs. C group could discriminate E group from C group with an area under curve (AUC) of 0.935 (training data) and 0.912 (testing data) (Fig. 5a). Moreover, the score formula for E vs. M group could also distinguish E group from M group with an AUC of 0.920 (training data) and 0.936 (testing data) (Fig. 5b).

**Fig. 5.**
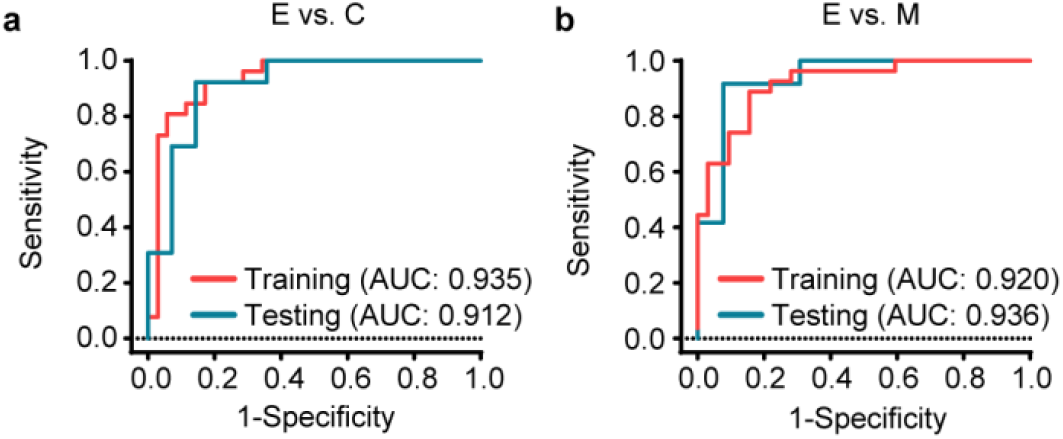
Receiver operating characteristic (ROC) curve analyses for the diagnostic performances of optimal gene signatures as predictors for occult tumor progression in mice. ROC curves for the diagnostic performances of the prediction score formulas generated from LASSO regression in the mouse cohort. ROC curves for the discrimination of early-early tumor (occult tumor, E group) from negative control group (C group) (**a**, E vs. C) or from macroscopic melanoma group (M group) (**b**, E vs. M). Probability statistics calculated according to the prediction score formulas generated from LASSO regression analyses: Probability = e^Score^ / (1 + e^Score^). 95% CI of AUC: training data 0.872-0.999, testing data 0.799-1.000 (**a**); training data 0.852-0.988, testing data 0.837-1.000 (**b**).

Although platelets have been suggested as a valuable platform for cancer diagnostics^3,20,21^, studies have yet to address their potential as a cancer screening platform. By using a mouse model that is simple, affordable and efficient, we identified differentially expressed RNAs in platelet signatures of mice injected with a suboptimal number of tumor cells, compared with mice with large melanomas and negative controls. These genes presented strong positive correlations with RNAs implicated in immune response and regulation. This possibly reflects the interactions between tumor cells and the immune system in the early stage of tumorigenesis. Moreover, the lack of enriched biological pathways from PBMC samples suggests platelets are the optimum biosource for early detection of cancer.

Indeed there are previous longitudinal studies using pre-diagnostic serums to screen for novel biomarkers of early cancer detection. However, these studies utilized pre-diagnostic serum to detect tumor-specific antigens or auto-antibodies for cancer risk prediction with limited sensitivity^2,22–24^. Novel cancer markers such as circulating tumor cells (CTCs) and circulating tumor DNA (ctDNA) offer new genomic approaches to screen for cancer through liquid biopsies. However, recent studies indicate CTCs assay cannot differentiate between patients with early-stage malignancy and people with no cancer and it has limited specificity as a screening tool^1,4^. On the other hand, ctDNA has promised to be a sensitive and specific test for cancer screening^25,26^. Still, ctDNA testing has several limitations for a screening platform compared with platelet RNA testing. First, the quantity of ctDNA is very limited even in cancer patients, not to mention in patients with early-stage cancer, while blood platelets are quite abundant. So the volume of blood needed in platelet testing is about 0.1 ml while ctDNA testing requires at least 10 ml. Second, the isolation and conversion process may cause damages to ctDNA, while platelet isolation procedure is simple and sample is stable and easy for storage. Third, ctDNA analysis could only detect frequently mutated genes in common cancers. The evolutionary and heterogeneity nature of cancer demands a large amount of possible mutations to be screened to achieve a consistent biomarker. Platelet biomarkers, on the other hand, are genes correlated with immune response and regulation according to our findings. Hence, platelet RNA testing may not be affected by cancer type or heterogeneity. Fourth, ctDNA extraction requires an expensive kit while platelet isolation needs no expensive consumables. The subsequent sequencing analysis of ctDNA is also more expensive than platelet sequencing in our study. Furthermore, our study used LASSO regression to select the optimum gene-expression-signature for the prediction of cancer risk via quantitative real-time PCR. Hence our strategy with the prediction models including 4 or 5 biomarkers as variables is much more cost-effective than ctDNA testing for hundreds of hotspots. Fifth, our platelet RNA prediction model could discriminate early-stage cancer from both healthy control and macroscopic tumor group, while biomarkers or screening models from previous studies often cannot distinguish samples from different stages of cancer. Thus platelet RNA testing may easily determine the best time for possible intervention. Last but not the least, platelet RNA testing described in our proof-of-concept study takes hours via qPCR while ctDNA testing takes days or weeks via next-generation sequencing and require skilled biology and bioinformatics technicians. Hence platelet testing is much less time-consuming and requires less training of technicians. This demonstrates the potential of platelets as a non-invasive screening platform for the detection of occult cancer.

The sensitivity and specificity of our model could further improve by including more samples or increasing RNA quantities to avoid invalid qPCR results from low-abundant genes, or by employing machine learning of large sequencing data for validation. Since it has been shown that platelet profiles are influenced by tumor type^3^, it is feasible to add tumor type markers into the gene-panel to determine the origin of cancer. It would also be interesting to investigate platelet profiles in immunoediting animal models to further understand the role of platelets in cancer-immune interactions. Combined, our study provides evidence for potential clinical relevance of blood platelets as a platform for liquid biopsy-based early detection of cancer.

## Methods

### Mice

C57BL/6 mice were bred in the Laboratory Animal Center, Health Science Center, Xi’an Jiaotong University. All mice were female and aged between 6–8 weeks at the beginning of all experiments. Animal experiments were approved by the Animal Ethics Committee of Xi’an Jiaotong University. All animal experiments were set with a maximum endpoint at a tumor volume of 1,000 mm^3^.

### B16F10 cell line

B16F10 cells negative for mycoplasma contamination were cultured and passaged in RPMI-1640 medium containing 10% fetal calf serum (FCS) at 37 °C/5% CO_2_. For animal inoculations, cells were cultured in Dulbecco’s Modified Eagle’s Medium (DMEM) supplemented with 10% FCS and 5 µg/ml plasmocin prophylactic reagent (InvivoGen, Cat.: ant-mpp) at 37 °C/5% CO_2_ to induce better melanin production. B16F10 cell line was generously provided by Dr Hui Zhang from Institute of Human Virology, Sun Yat-sen University and originally purchased from the ATCC.

### B16F10 melanoma inoculation

For melanoma inoculation, B16F10 melanoma cells were harvested by washing with phosphate buffer saline (PBS), then incubating cells at 37 °C for 1–2 min with 1 × Trypsin/EDTA solution and washing with Hanks’ balanced saline solution (HBSS) twice. For B16F10 inoculation, the right flanks of mice were shaved with a mini-razor and cells (2 × 10^3^, 5 × 10^3^, 1 × 10^4^ and 1 × 10^5^ cells per mouse for each group) were suspended in 100 μl HBSS and then injected under the right flank subcutaneously using a 30G needle. Tumor formation was monitored by inspecting via a magnifying scope and measured with a caliper periodically (tumor volume was estimated using this formula: volume = length × width × height × 0.5)^27^.

### Blood sample collection

Blood samples were collected from retro-orbital sinuses of C57BL/6 mice 14 days after inoculation. For terminal or nonterminal blood collection, mice were fully anesthetized with isoflurane and blood samples were collected by puncturing the retro-orbital sinuses of mice using microhematocrit capillary tubes. Blood was collected into a tube containing the anti-coagulant EDTA. After nonterminal blood collection (less than 1% of body weight, approximately 150-200 μl), the tube was withdrawn and a slight pressure was put on the eye with a sterile cotton swab to ensure hemostasis. After terminal blood collection, mice were euthanized by cervical dislocation.

### Isolation of platelet and PBMC RNA

Blood platelets isolation was performed as described previously^3,28^. Briefly, anti-coagulated blood was centrifuged at 180 × g at room temperature for 10 min, yielding platelet-rich plasma. Platelets were isolated from the platelet-rich plasma by centrifugation at room temperature for 10 min at 1,250 × g. The platelet pellet was lysed in TRIzol Reagent (Invitrogen, Thermofisher Scientific) and frozen at −80 °C for future use.

The bottom layer of the centrifuged blood sample from the first step of platelet isolation was further used for peripheral blood mononuclear cell (PBMC) isolation using mouse PBMC isolation kit (TBD science, Tianjin) following the manufacturer’s instructions. Briefly, the aforementioned bottom layer was mixed with the same volume of diluting solution provided by the manufacturer and the mixture was carefully layered on the PBMC isolation reagent of the same volume in a sterile centrifuge tube. The tube was then centrifuged at 950 × g for 30 min at room temperature. PBMC layer was transferred into a new tube from the interphase with a transfer pipette and washed twice by mixing with 10 ml washing solution (also provided by the manufacturer) followed by centrifuging at 250 × g for 10 min. The final PBMC pellet was lysed in TRIzol and kept in −80 °C for further use.

### Next generation sequencing

Next generation sequencing was performed in Novogene (Tianjin, China). Briefly, platelet or PBMC RNA samples were assessed for quantity, purity and integrity using NanoPhotometer® spectrophotometer (IMPLEN, CA, USA) and RNA Nano 6000 Assay Kit of Bioanalyzer 2100 system (Agilent Technologies, CA, USA). Samples were pooled to satisfy the quantity criteria of RNA sequencing and a minimum amount of 20 ng RNA per pooled sample was used as input material for the RNA sample preparations. For platelet samples, the numbers of mice sacrificed for 5 pooled samples in three groups were as follows: 3, 4, 5, 3 and 5 mice for O group (optimal inoculation group, mice inoculated with 1 × 10^5^ B16F10 cells); 10, 10, 9, 11 and 10 mice for S group (suboptimal inoculation group, mice inoculated with 2 × 10^3^ B16F10 cells); 10, 5, 5, 10 and 10 mice for C group (negative control group, mice injected with HBSS). For PBMC samples, the numbers of mice sacrificed for 5 pooled samples in three groups were as follows: 3, 4, 5, 3 and 5 mice for O group; 5, 4, 5, 5 and 5 mice for S group; 5 mice for each sample in C group.

Sequencing libraries were generated using NEBNext^®^ UltraTM RNA Library Prep Kit for Illumina^®^ (NEB, USA) following the manufacturer’s recommendations with index codes added to attribute sequences to each sample. Briefly, mRNA was purified from total RNA using poly-T oligo-attached magnetic beads. Fragmentation of mRNA was carried out followed by cDNA synthesis. Sequencing libraries were created by converting RNA to cDNA via reverse transcription and adding specialized adapters to both ends. Next, library fragments were purified and then PCR was performed with Phusion High-Fidelity DNA polymerase. PCR products were purified (AMPure XP system) and library quality was assessed on the Agilent Bioanalyzer 2100 system (Agilent Technologies, CA, USA). Finally, qualified libraries were sequenced on an Illumina Novaseq platform.

### Sequencing data analysis

Novogene provided sequence alignment, data mapping and differential expression analysis. Briefly, raw data of fastq format were processed to obtain clean reads with high quality by removing reads containing adapter, reads containing ploy-N and reads of low quality from raw data. Clean reads were aligned to mus musculus reference genome using Hisat2 v2.0.5 alignment program. The R package featureCounts v1.5.0-p3 was applied to count the reads numbers mapped to each gene^29^. And then FPKM (fragments per kilobase of transcript sequence per millions base pairs sequenced) of each gene was calculated based on the length of the gene and read count mapped to this gene.

Differential expression analysis was performed using DESeq2 R package 1.16.1^30^. The resulting *P* values were adjusted using the Benjamini and Hochberg’s approach for controlling the false discovery rates. Genes with *P* values < 0.05 were assigned as differentially expressed for pairwise comparisons. Differentially expressed genes for subsequent screening of distinct markers of early-early cancer (eDEGs) must meet these requirements: log_2_(S vs. C Fold Change) > 0, *P* < 0.05, log_2_(S vs. O Fold Change) > 0, *P* < 0.05; or log_2_(S vs. C Fold Change) < 0, *P* < 0.05, log_2_(S vs. O Fold Change) < 0, *P* < 0.05. O group, optimal inoculation group, mice inoculated with 1 × 10^5^ B16F10 cells; S group, suboptimal inoculation group, mice inoculated with 2 × 10^3^ B16F10 cells; C group, negative control group, mice injected with HBSS. Bioinformatics analyses and data visualizations.

Data visualizations were performed using R (version 3.6.3)^31^. Heatmaps and clusterings were generated using pheatmap package^32^. Dot plots and bubble plots were generated using ggplot2 and corrplot packages^33,34^. Pathway enrichment analyses of differentially expressed genes in suboptimal inoculation group (eDEGs) were performed using clusterProfiler package with reference from KEGG (Kyoto Encyclopedia of Genes and Genomes) pathways with *P* values adjusted by Benjamini and Hochberg method^35,36^. The protein-protein interaction (PPI) network of eDEGs was retrieved from STRING database^18^ and reconstructed using Cytoscape^37^. Each node’s degree of connectivity in the network was calculated. Molecular COmplex DEtection (MCODE)^38^ was used to find gene clusters based on topology locating densely connected regions.

### Quantitative real-time PCR

Blood platelet RNA was isolated as described above. Platelet RNA was then converted to complementary cDNA using PrimeScript RT Master Mix (Takara) according to the manufacturer’s instructions. Quantitative real-time PCR (qPCR) was performed with TB Green Premix Ex Taq (Takara) using LightCycler 96 System (Roche Life Science) with parameters adjusted according to the PCR cycler and the enzyme’s manuals. The reaction process was as follows: preincubation at 95 °C for 30 s; 40 cycles of 5 s at 95 °C and 30 s at 55 °C for annealing and extension; melting at 95 °C for 5 s, 60 °C for 60 s and 95 °C for 1 s; cooling for 30 s at 50 °C. Cycle-threshold (Ct) values were determined for each gene and normalized to the housekeeping genes *Actb, Gapdh, Gusb, Gnas* and *Oaz1*, which were validated previously as reliable reference genes for platelet RNA qPCR^39,40^.

### Statistic analyses

All statistical analyses were performed in SPSS 18.0 or Prism 8 (Graphpad) and were two-sided. All experiments were performed with replicates as indicated, either with representative data shown, or with pooled data shown. Figures with pooled data from multiple experiments included all experiments performed. All data reflected multiple independent experiments with at least 3 mice per experiment, in which similar results were obtained.

Kaplan-Meier curves were generated to illustrate the relationship between percentage of tumor-free mice and time after inoculation. Mantel-Cox tests were used to test statistic significance.

Joinpoint software version 4.8.0.1 was used to analyze tumor growth data for multi-phase regression in order to determine when the tumor went from occult state into fast progressing phase^19^. A maximum of 1 joinpoint was allowed based on the number of data points and previous studies on the role of angiogenesis in tumor dormancy^41^. The statistic significance of the change in tumor growth trend over time was tested using a Monte Carlo Permutation method embedded in the Joinpoint software^19^. Blood samples from mice whose tumor became visible more than 5 days after blood collection (19 days post-inoculation) were categorized as early-early group (E group). Alternatively, samples from mice with a mini-tumor (tumor volume < 1 mm^3^) on the day of blood collection (14 days post-inoculation) that did not progress for at least 20 days since inoculation (joinpoint ≥ 20, Permutation test *P* value < 0.05) were also classified as early-early group (E group). Blood samples from mice with macroscopic and palpable tumors (tumor volume > 30 mm^3^) were categorized as melanoma group (M group) and blood samples from mice injected with HBSS were in negative control group (C group).

Statistic analyses of qPCR results were performed via SPSS 18.0. Kruskal-Wallis non-parametric test was executed and adjusted *P* value below 0.05 was assigned as significant. Samples with more than 10 genes with invalid Ct values (no signal within 40 cycles of PCR due to low RNA quantity) out of 36 markers were excluded. Then genes with invalid Ct values in at least 5 samples in both dependent variable groups (probably due to low expression levels of certain chosen markers in platelets) were not included for subsequent variable selection via LASSO binary regression analysis.

The Least Absolute Shrinkage and Selection Operator (LASSO) model is a shrinkage method for regression with high-dimensional predictors, which can preserve valuable variables from a large and potentially multicollinear set of variables, and avoid overfitting. This method is suited for analyzing gene expression data where multicollinearity of selected genes in related biological pathways may occur. We performed LASSO binary logistic regression using glmnet R package^42^. Data were randomly divided into training set (70%) and testing set (30%). We utilized ten-fold cross-validation to select the penalty term λ. The binomial deviance was set as measures of the predictive performance of the fitted models. The built-in function in glmnet package produced the λ that minimized the binomial deviance. The coefficients of selected variables were obtained through the penalizing process. The seed was set to 10 for data replication. The prediction score formulas for the discrimination of early-early tumor (occult tumor, E group) from negative control group (C group) or from macroscopic melanoma group (M group) were established as follows: Score = Intercept + S Coefficient × (Ct_Variable_ – Ct_Ref_).

Receiver operating characteristic (ROC) curves were constructed and area under curve (AUC) was calculated using SPSS 18.0. Probability statistics for ROC were calculated according to the prediction score formulas generated from LASSO regression analyses described above: Probability = e^Score^ / (1 + e^Score^).

### Data availability

The accession number for the raw sequencing data reported in this study is GEO: GSE160650.

## Acknowledgements

Financial support was provided by Xinjiang Guan Run Logistics Co., Ltd and J.R.. (Funding No.: 20181224) and Ruilan Jiang. A special thank you goes to Dr. Le Ma, Dr. Liang Bai, Dr. Bei Han, Dr. Jing Han, Dr. Jing Lin, Dr. Weiqing Wang, Yixue Li, Qiao Peng, Jianing Wu and Ruilei Cheng for their help in this study.

## Author contributions

Y.Y. and J.R. designed the study and performed the experiments. L.Z., Y.N., F.J., J.H., L.J., S.L, W.L., L.X., Z.K., and C.D. participated in animal maintenance, animal blood sample collection and/or tumor cell inoculation. Y.Y. and S.M. performed data analyses and interpretation. Y.Y. wrote the manuscript. All authors approved the final manuscript.

## Figures, extended data figures and tables, and supplementary information

**Extended Data Fig. 1.**
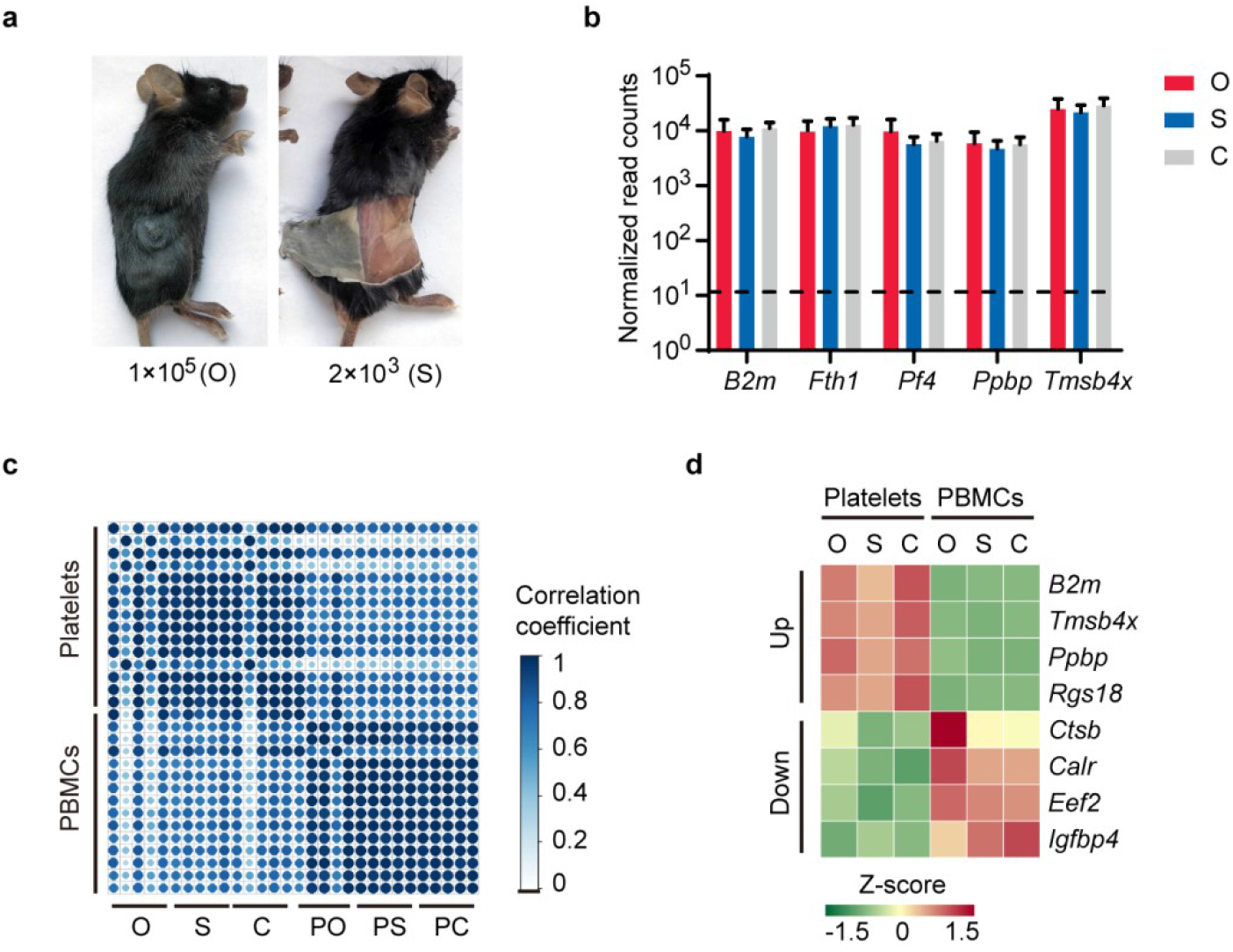
Platelet RNA profiles of mice inoculated with B16F10 cells are consistent with platelet signatures in previous studies. **a**, Images of mice after terminal blood collection. Representative images of mice in O group (optimal inoculation group, mice inoculated with 1 × 10^5^ B16F10 cells) (left) and S group (suboptimal inoculation group, mice inoculated with 2 × 10^3^ B16F10 cells) (right). **b**, Platelet mRNA sequencing data of known platelet-abundant genes with a dashed line showing the average read count of our data. **c**, Pearson’s correlation (color bar) matrix of our mRNA sequencing data of platelets and PBMCs (columns and rows). S: platelet suboptimal inoculation group; O: platelet optimal inoculation group; C: platelet negative control group. PS: PBMC suboptimal inoculation group; PO: PBMC optimal inoculation group; PC: PBMC negative control group. **d**, Heatmap of previously reported differentially expressed genes between platelets and PBMCs from our sequencing data.

**Extended Data Fig. 2.**
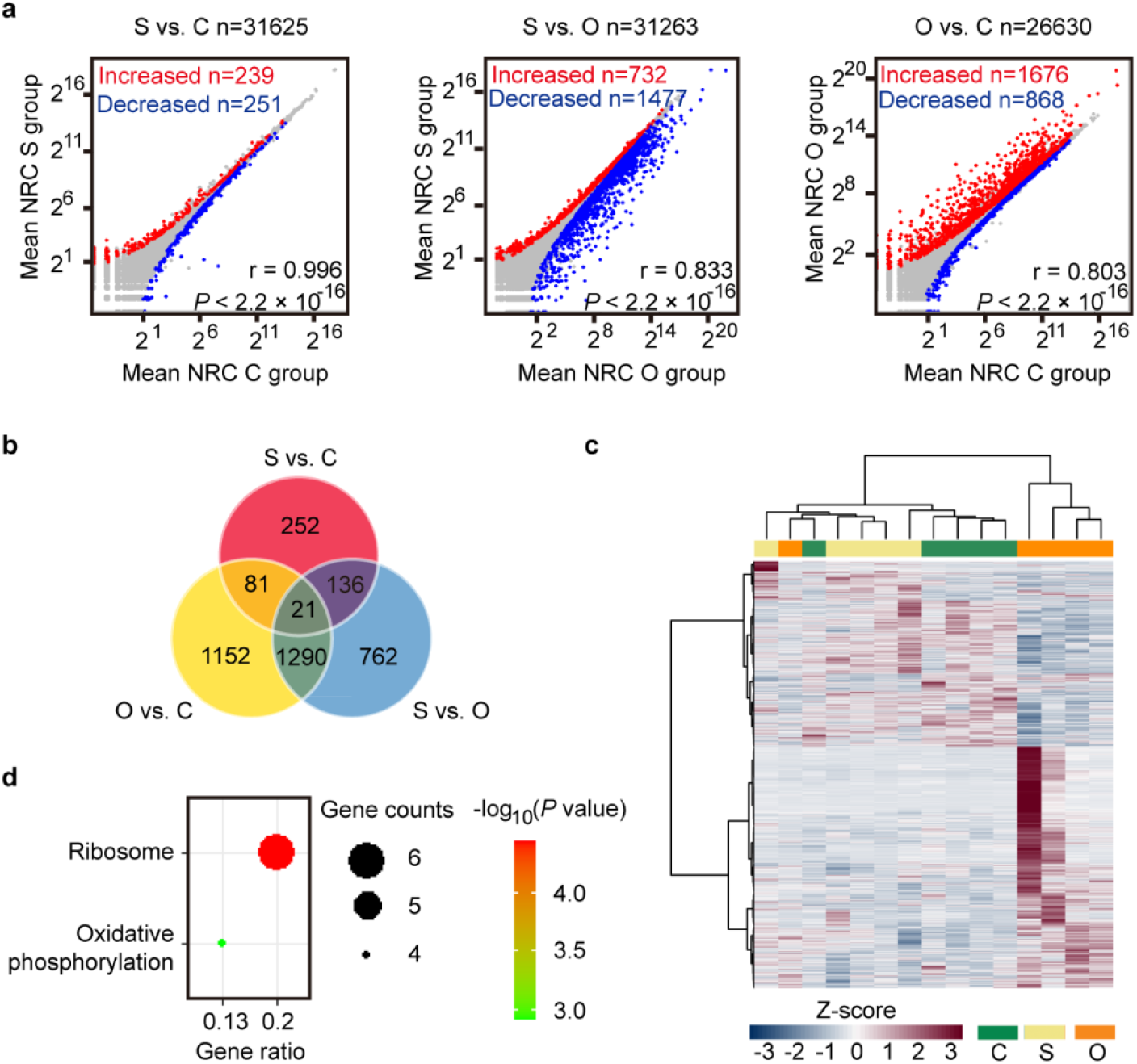
PBMC RNA profiles of mice inoculated with an optimal and a suboptimal number of B16F10 cells. **a**, Correlation plots of mRNAs detected in PBMCs of suboptimal inoculation group (S group, mice inoculated with 2 × 10^3^ B16F10 cells), negative control group (C group, mice injected with HBSS) and optimal inoculation group (O group, mice inoculated with 1 × 10^5^ B16F10 cells) mice, including highlighted increased (red) and decreased (blue) PBMC mRNAs. NRC, normalized read counts (mean of group). r value calculated from Pearson’s correlation test. **b**, Venn diagram of differentially expressed genes from pairwise comparisons. **c**, Heatmap of hierarchical clustering of PBMC mRNA profiles of S group (beige), C group (green) and O group (orange). Data pooled from (**a, b**) *n* = 5 biologically independent experiments with *n* = 24 (S group) or *n* = 25 (C group) and *n* = 20 (O group) mice or data representing all 5 independent experiments (**c**). **d**, Top GO terms of pathway enrichment analysis of eligible eDEGs (strategy same as Fig. 3a, details in “Methods”) in PBMCs with reference from KEGG pathways. Adjusted *P* value < 0.05, Benjamini and Hochberg method.

**Extended Data Fig. 3.**
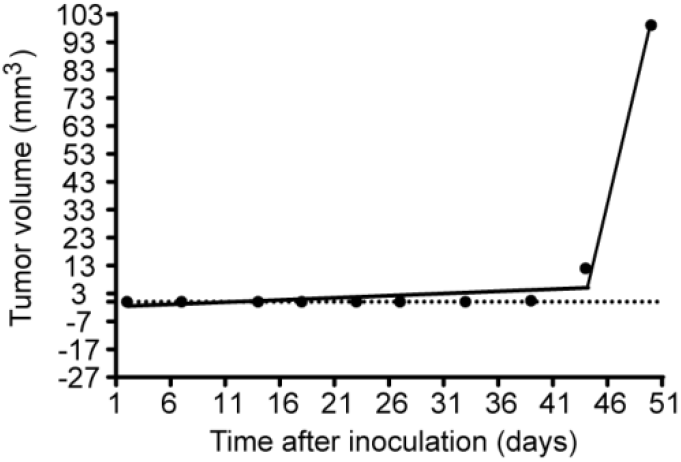
An example of the selected models from Joinpoint multi-phase regression analyses. Tumor growth data from one mouse in early-early tumor group (E group).

**Extended Data Fig. 4.**
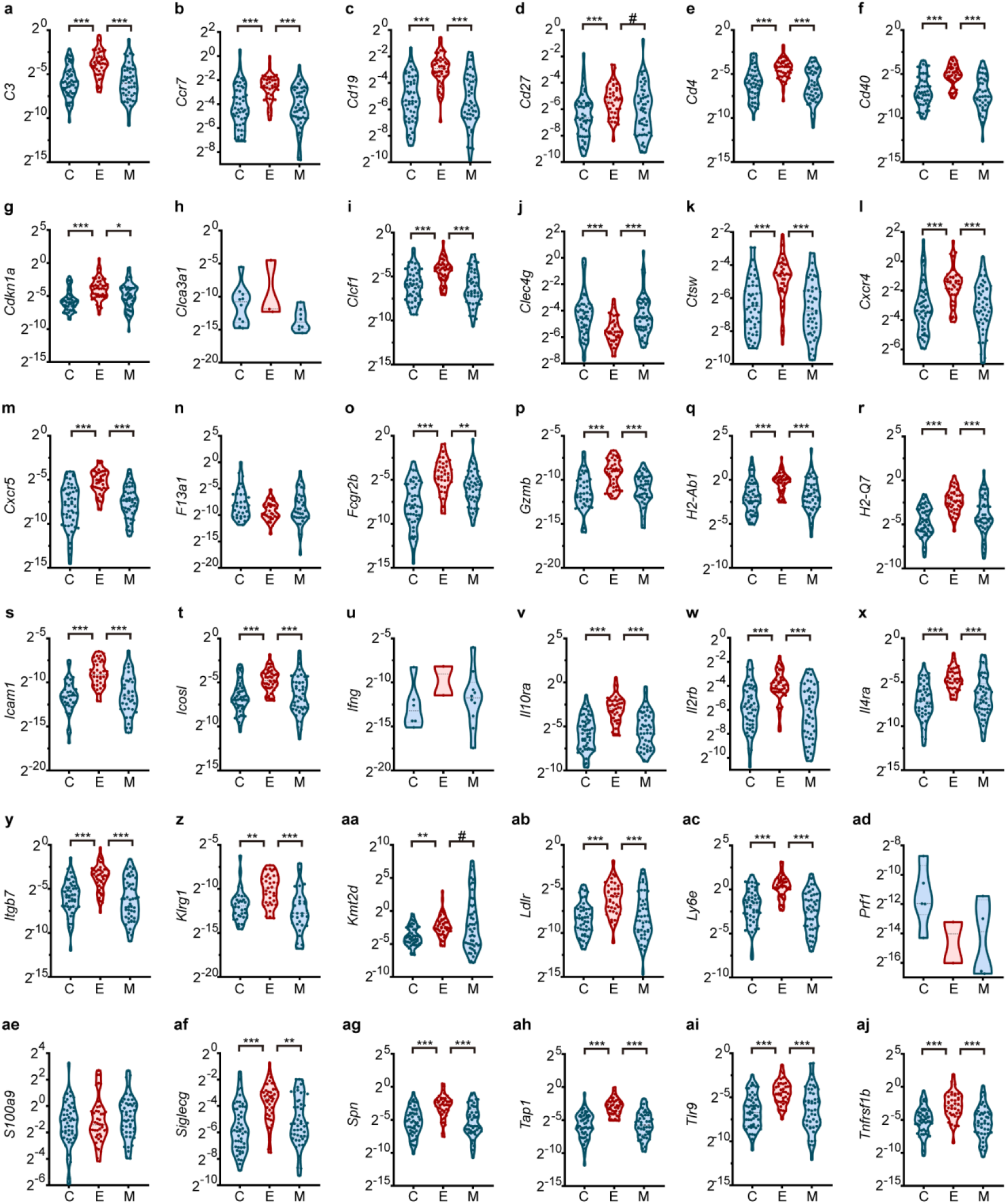
Quantitative real-time PCR verifications of the expression levels of selected 36 genes in the mouse cohort. Violin plots of normalized gene expression levels (2^ΔCt(Ref-Gene)^) of 36 selected genes in three groups of the mouse cohort. C: negative control; E: early-early tumor; M: macroscopic melanoma. \**P* < 0.05, \*\**P* < 0.01, \*\*\**P* < 0.001, Kruskall-Wallis test. Data without significance tags representing non-significant for analyses between three groups (**h, n, u, ad, ae**) (detailed statistics including *n* values and *P* values see Extended Data Table 5).

**Extended Data Fig. 5.**
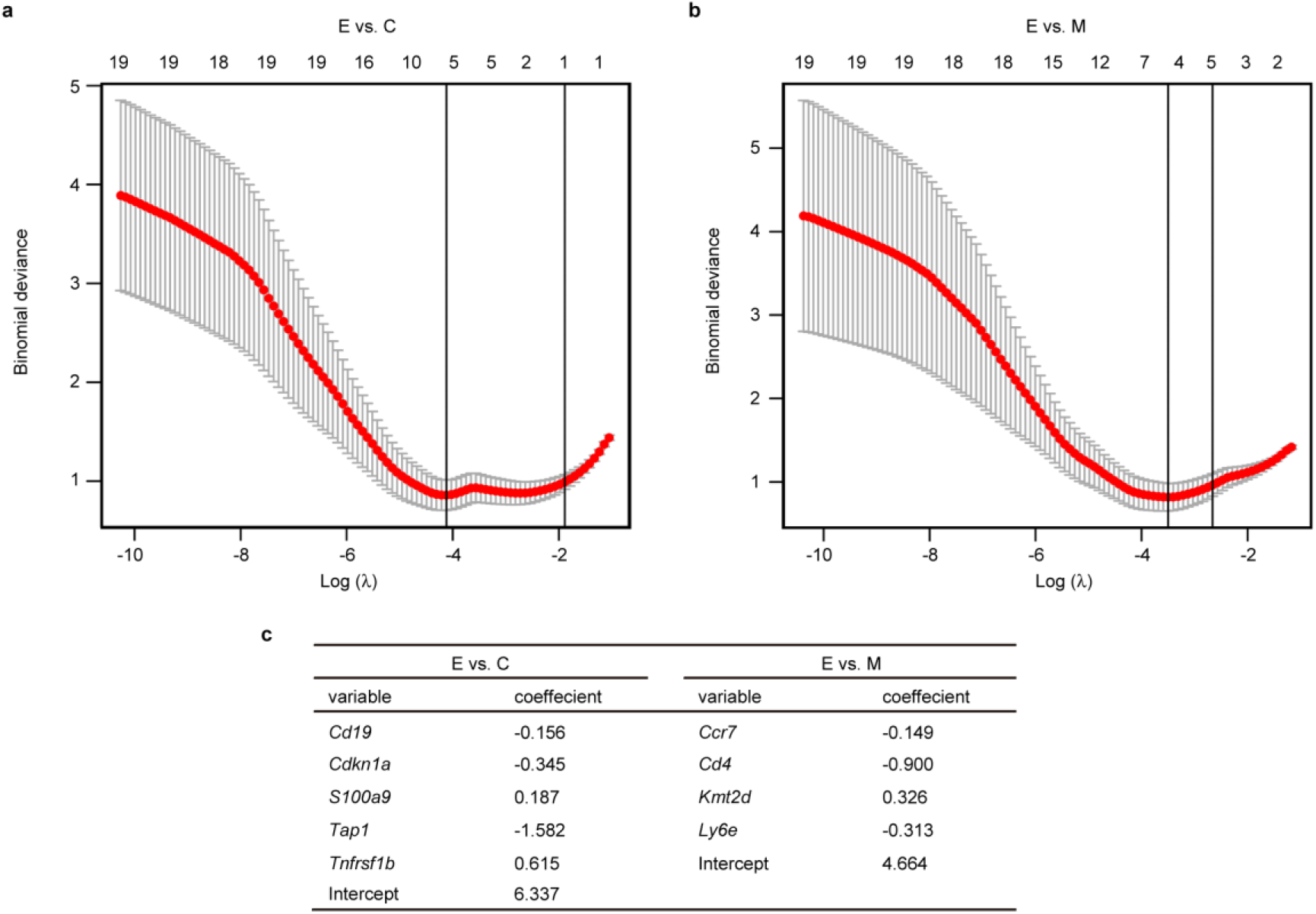
LASSO regression model construction and variable selection for predicting occult tumor progression in the mouse cohort. **a, b**, Ten-fold cross-validation for the selection of the penalty term λ with the binomial deviance as measures of the predictive performance of the fitted models. The dependent variable groups: E vs. C (**a**) or E vs. M (**b**). **c**, Coefficients derived from LASSO regression. E, early-early tumor (occult tumor that progressed into macroscopic tumor later); C, negative control group; M, macroscopic melanoma group. Numbers of samples included in LASSO regression: E, *n* = 40; C, *n* = 50; M, *n* = 45. Numbers of variables included in LASSO regression: E vs. C, *n* = 29; E vs. M, *n* = 30. The prediction score formulas for the discrimination of E group from C group (Score_EC_) or from M group (Score_EM_) established as follows: Score_EC_ = 6.337 – 0.156 × (Ct_*Cd19*_ – Ct_Ref_) – 0.345 × (Ct_*Cdkn1a*_ – Ct_Ref_) + 0.187 × (Ct_*S100a9*_ – Ct_Ref_) – 1.582 × (Ct_*Tap1*_ – Ct_Ref_) + 0.615 × (Ct_*Tnfrsf1b*_ – Ct_Ref_); Score_EM_ = 4.664 – 0.149 × (Ct_*Ccr7*_ – Ct_Ref_) – 0.900 × (Ct_*Cd4*_ – Ct_Ref_) + 0.326 × (Ct_*Kmt2d*_ – Ct_Ref_) – 0.313 × (Ct_*Ly6e*_ – Ct_Ref_).

**Extended Data Table 1.** Enriched KEGG pathways of differentially expressed mRNAs in platelets of suboptimal inoculation group compared with optimal inoculation group and control group.

| Term | Count | Gene ratio | *P* value | *P* adjust |
| --- | --- | --- | --- | --- |
| Cytokine-cytokine receptor interaction | 24 | 0.110 | 0.000 | 0.000 |
| Cell adhesion molecules | 20 | 0.091 | 0.000 | 0.000 |
| Epstein-Barr virus infection | 18 | 0.082 | 0.000 | 0.000 |
| Human T-cell leukemia virus 1 infection | 18 | 0.082 | 0.000 | 0.001 |
| Th1 and Th2 cell differentiation | 14 | 0.064 | 0.000 | 0.000 |
| Th17 cell differentiation | 14 | 0.064 | 0.000 | 0.000 |
| Phagosome | 14 | 0.064 | 0.000 | 0.002 |
| Human immunodeficiency virus 1 infection | 14 | 0.064 | 0.002 | 0.022 |
| Hematopoietic cell lineage | 12 | 0.055 | 0.000 | 0.000 |
| Staphylococcus aureus infection | 11 | 0.050 | 0.000 | 0.003 |
| Lysosome | 11 | 0.050 | 0.000 | 0.004 |
| Tuberculosis | 11 | 0.050 | 0.005 | 0.039 |
| Allograft rejection | 10 | 0.046 | 0.000 | 0.000 |
| Rheumatoid arthritis | 10 | 0.046 | 0.000 | 0.001 |
| Viral myocarditis | 10 | 0.046 | 0.000 | 0.001 |
| Antigen processing and presentation | 10 | 0.046 | 0.000 | 0.001 |
| Systemic lupus erythematosus | 10 | 0.046 | 0.004 | 0.030 |
| Primary immunodeficiency | 9 | 0.041 | 0.000 | 0.000 |
| Autoimmune thyroid disease | 9 | 0.041 | 0.000 | 0.002 |
| Viral protein interaction with cytokine and cytokine receptor | 9 | 0.041 | 0.001 | 0.006 |
| Chagas disease | 9 | 0.041 | 0.001 | 0.009 |
| Intestinal immune network for IgA production | 8 | 0.037 | 0.000 | 0.000 |
| Inflammatory bowel disease | 8 | 0.037 | 0.000 | 0.002 |
| Graft-versus-host disease | 8 | 0.037 | 0.000 | 0.002 |
| Type I diabetes mellitus | 8 | 0.037 | 0.000 | 0.004 |
| B cell receptor signaling pathway | 8 | 0.037 | 0.001 | 0.008 |
| PD-L1 expression and PD-1 checkpoint pathway in cancer | 8 | 0.037 | 0.001 | 0.013 |
| T cell receptor signaling pathway | 8 | 0.037 | 0.004 | 0.031 |
| Malaria | 6 | 0.027 | 0.003 | 0.023 |
| Glycosaminoglycan biosynthesis - chondroitin sulfate / dermatan sulfate | 5 | 0.023 | 0.000 | 0.002 |
| Asthma | 5 | 0.023 | 0.000 | 0.004 |

**Extended Data Table 2.** Enriched KEGG pathways of differentially expressed mRNAs in PBMCs of suboptimal inoculation group compared with optimal inoculation group and control group.

| Description | Count | Gene ratio | *P* value | *P* adjust |
| --- | --- | --- | --- | --- |
| Ribosome | 6 | 0.200 | 0.000 | 0.002 |
| Oxidative phosphorylation | 4 | 0.133 | 0.001 | 0.044 |

**Extended Data Table 3.** Joinpoint multi-phase regression statistics for all mice in early-early group of the mouse cohort.

| Mice | Joinpoint | *P* value |
| --- | --- | --- |
| E1 | 34 | 0.024 |
| E2 | 29 | 0.036 |
| E3 | 34 | 0.012 |
| E4 | 34 | 0.019 |
| E5 | 34 | 0.045 |
| E6 | 28 | 0.003 |
| E7 | 23 | 0.000 |
| E8 | 23 | 0.007 |
| E9 | 38 | 0.000 |
| E10 | 19 | 0.007 |
| E11 | 28 | 0.000 |
| E12 | 28 | 0.000 |
| E13 | 44 | 0.000 |
| E14 | 33 | 0.000 |
| E15 | 27 | 0.000 |
| E16 | 27 | 0.000 |
| E17 | 27 | 0.000 |
| E18 | 33 | 0.000 |
| E19 | 27 | 0.000 |
| E20 | 27 | 0.000 |
| E21 | 23 | 0.000 |
| E22 | 27 | 0.000 |
| E23 | 21 | 0.003 |
| E24 | 21 | 0.000 |
| E25 | 31 | 0.000 |
| E26 | 23 | 0.003 |
| E27 | 23 | 0.029 |
| E28 | 23 | 0.000 |
| E29 | 29 | 0.000 |
| E30 | 23 | 0.003 |
| E31 | 23 | 0.003 |
| E32 | 35 | 0.000 |
| E33 | 25 | 0.000 |
| E34 | 25 | 0.003 |
| E35 | 20 | 0.000 |
| E36 | 25 | 0.011 |
| E37 | 25 | 0.005 |
| E38 | 25 | 0.000 |
| E39 | 20 | 0.000 |
| E40 | 25 | 0.000 |
| E41 | 25 | 0.004 |
| E42 | 20 | 0.018 |
| E43 | 25 | 0.015 |
| E44 | 25 | 0.015 |

*P* value, the statistic significance of the change in tumor growth trend over time calculated via a Monte Carlo Permutation method

**Extended Data Table 4.**
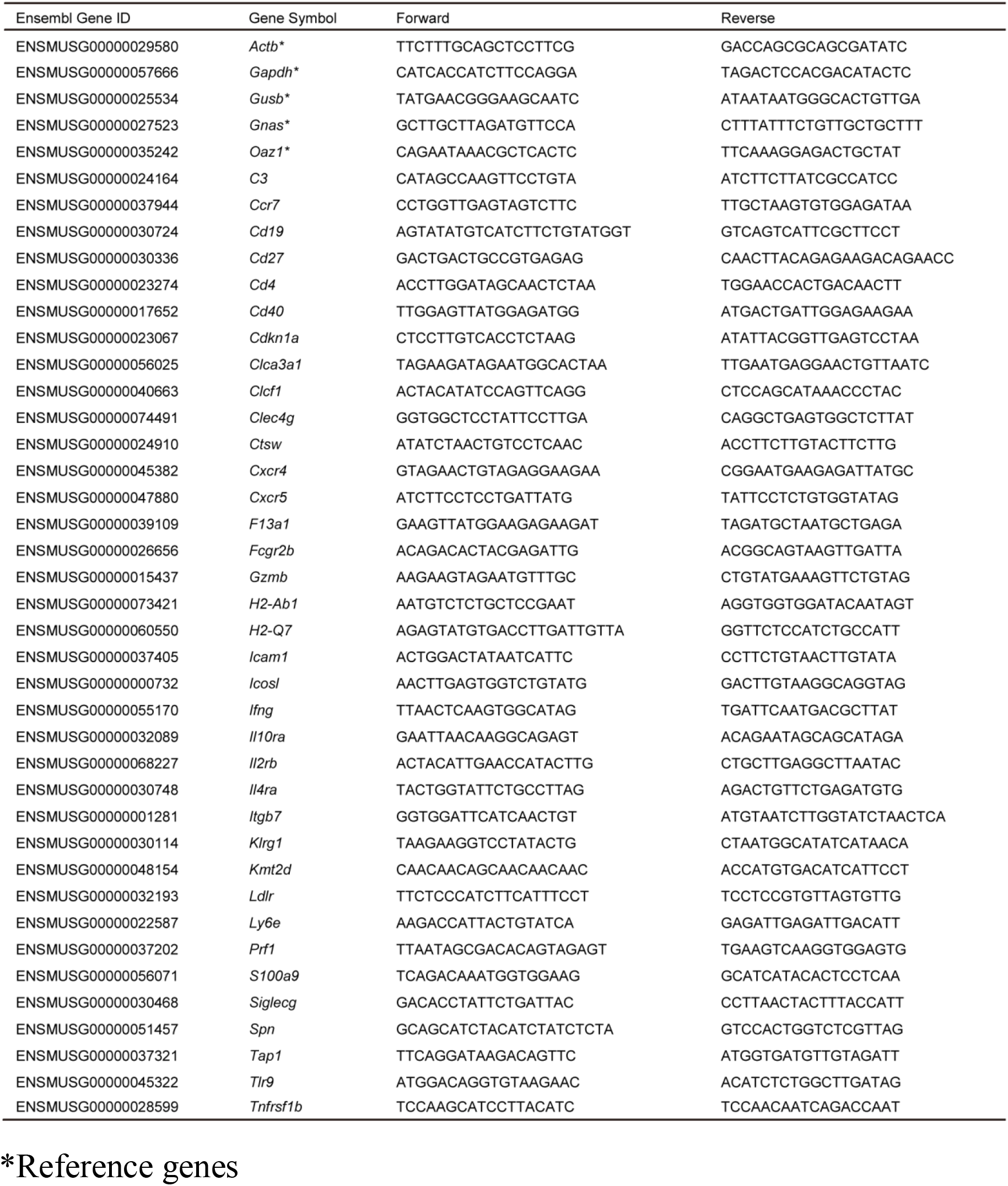
Primers for qPCR experiments.

**Extended Data Table 5.**
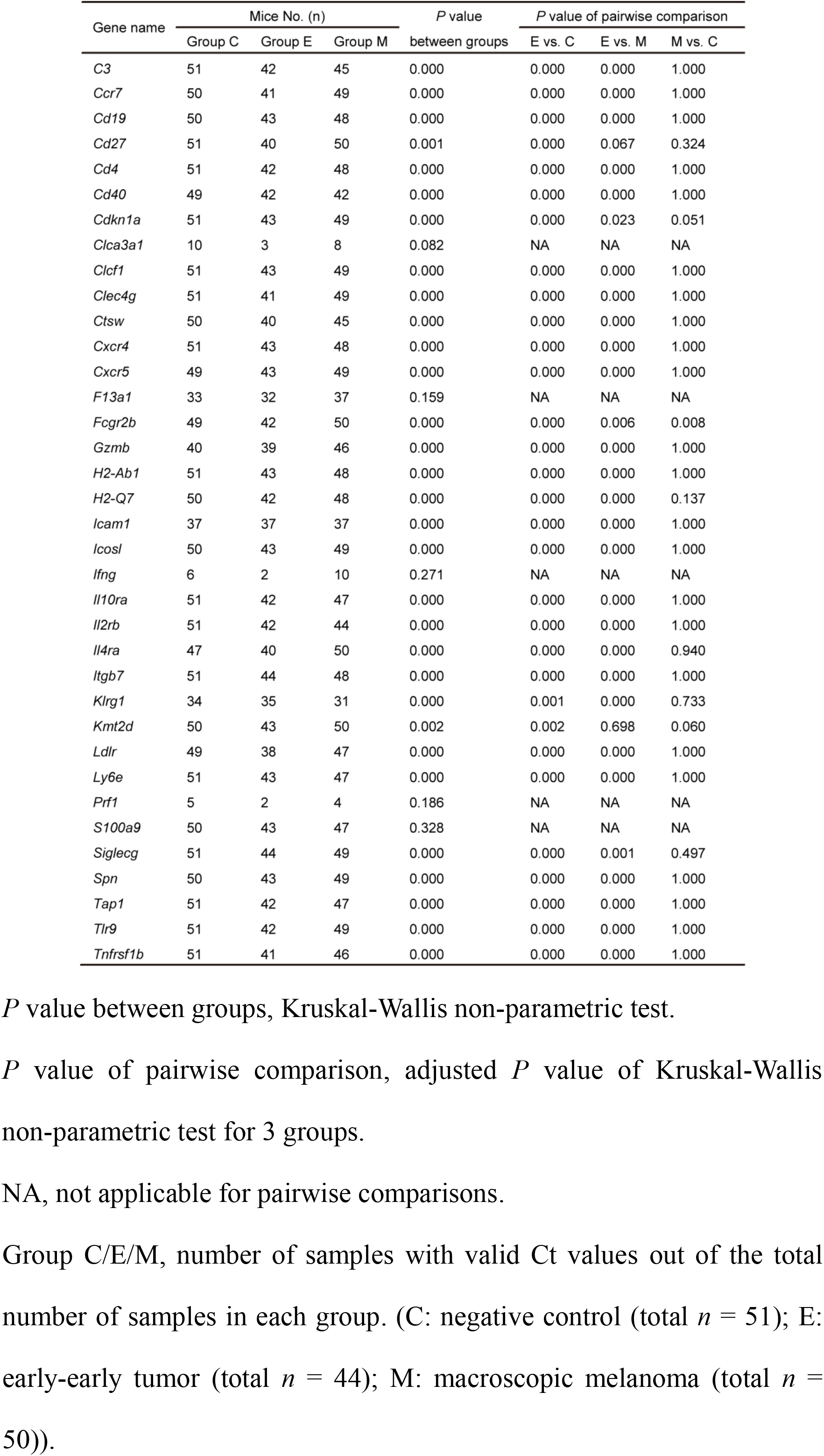
Statistic analyses of the normalized expression levels of selected 36 genes in the mouse cohort.

### Additional information

Supplementary Table 1 Differentially expressed mRNAs in platelets of suboptimal inoculation group compared with optimal inoculation group and control group.

Supplementary Table 2 Differentially expressed mRNAs in PBMCs of suboptimal inoculation group compared with optimal inoculation group and control group.

